# Pancreatic beta-cell specific deletion of VPS41 causes diabetes due to defects in insulin secretion

**DOI:** 10.1101/2020.04.25.043729

**Authors:** Christian H. Burns, Belinda Yau, Anjelica Rodriguez, Jenna Triplett, Drew Maslar, You Sun An, Reini E.N. van der Welle, Ross G. Kossina, Max R. Fisher, Gregory W. Strout, Peter O. Bayguinov, Tineke Veenendaal, David Chitayat, James A.J. Fitzpatrick, Judith Klumperman, Melkam A. Kebede, Cedric S. Asensio

## Abstract

Insulin secretory granules (SGs) mediate the regulated secretion of insulin, which is essential for glucose homeostasis. The basic machinery responsible for this regulated exocytosis consists of specific proteins present both at the plasma membrane and on insulin SGs. The protein composition of insulin SGs thus dictates their release properties, yet the mechanisms controlling insulin SG formation, which determines this molecular composition, remain poorly understood. VPS41, a component of the endo-lysosomal tethering HOPS complex, was recently identified as a cytosolic factor involved in the formation of neuroendocrine and neuronal granules. We now find that VPS41 is required for insulin SG biogenesis and regulated insulin secretion. Loss of VPS41 in pancreatic β-cells leads to a reduction in insulin SG number, changes in their transmembrane protein composition, and defects in granule regulated exocytosis. Exploring a human point mutation, identified in patients with neurological but no endocrine defects, we show that the effect on SG formation is independent of HOPS complex formation. Finally, we report that mice with a deletion of VPS41 specifically in β-cells develop diabetes due to severe depletion of insulin SG content and a defect in insulin secretion. In sum, our data demonstrate that VPS41 contributes to glucose homeostasis and metabolism.

## Introduction

Proper temporal release of peptide hormones plays a critical role in the body’s ability to maintain homeostasis. Pancreatic β-cells, for example, produce insulin and store it within intracellular vesicles, known as insulin secretory granules (SGs). At any given time, these cells can accumulate thousands of insulin SGs, and these vesicles undergo exocytosis only in response to extracellular stimuli (e.g., hyperglycemia). The basic machinery responsible for this regulated exocytosis consists of proteins present both at the plasma membrane and on insulin SGs. Thus, the protein composition of insulin SGs partly dictates their release properties. SGs form at the trans-Golgi network (TGN), where they bud as immature granules. These vesicles then undergo a series of maturation steps, which includes lumenal acidification, processing of proinsulin to insulin, insulin crystallization with zinc and removal of immature granule markers (e.g., syntaxin 6, VAMP4) (Ahras et al., 2006; Borgonovo et al., 2006; Dittie et al., 1996; Hou et al., 2009; Klumperman et al., 1998; Park et al., 2009; Takeuchi & Hosaka, 2008; Tooze, 1998). Surprisingly, the exact mechanisms dictating SG biogenesis and maturation, which is thought to determine their molecular composition, remains poorly understood.

VPS41 and VPS39 associate with the VPS Class C core proteins (VPS11, VPS16, VPS18 and VPS33A) to form the homotypic fusion and vacuole protein sorting (HOPS) complex, which acts as a tether for late endosome-lysosome fusion (Price et al., 2000; Rieder & Emr, 1997; Seals et al., 2000; van der Beek et al., 2019). Disruption in HOPS impairs delivery of endocytic cargo to lysosome (Balderhaar & Ungermann, 2013) and causes defects in autophagosome clearance (Jiang et al., 2014; Takats et al., 2014). Independently of its role in HOPS, VPS41 also functions in the fusion of TGN-derived LAMP carriers with late endosomes (Pols et al., 2013). Interestingly, gene expression analysis revealed a significant reduction in VPS41 mRNA levels in islets of a model of mice susceptible to develop diabetes (Keller et al., 2008). Beside this, several VPS41 variants have been associated with type 2 diabetes in humans, suggesting that it might be a factor contributing to the development of the disease (Type_2_Diabetes_Knowledge_Portal). Furthermore, we previously found that VPS41 affects regulated secretion of neuropeptides from neuroendocrine cells and neurons, in a process that is independent of the HOPS complex (Asensio et al., 2013). Altogether, this led us to investigate if VPS41 contributes to glucose homeostasis by influencing insulin SG biology.

Using CRISPR/Cas9 in rat insulinoma INS-1 cells, we now show that VPS41 is required for proper insulin storage and regulated secretion. Lack of VPS41 decreases SG number and affects their ability to undergo regulated exocytosis, which is concomitant with a defect in sorting of SG transmembrane proteins. We further show that a patient mutation of VPS41, which is deficient for HOPS function (van der Welle et al., 2019), rescues insulin regulated secretion in VPS41 KO INS-1 cells, indicating VPS41 function in insulin regulated secretion is independent of the HOPS complex. Finally, we report that mice with a targeted deletion of VPS41 in β-cells develop diabetes due to a defect in insulin SG biogenesis and secretion. These studies for the first time implicate VPS41 in regulating β-cell function and control of glucose homeostasis.

## Results

### Loss of VPS41 decreases insulin regulated secretion and cellular content in INS-1 cells

To assess the potential contribution of VPS41 to insulin secretion, we generated VPS41 KO INS-1 cells using CRISPR/Cas9. The analysis of indels from genomic DNA isolated from these cells revealed the presence of two distinct single base pair insertions predicted to mutate the initiator codon for VPS41 (**Supplemental Figure 1A**). In addition, we could not detect VPS41 protein by western blot in lysates generated from these cells, even when using sensitive enhanced chemiluminescence reagents (**Figure 1A**). As a control for potential Cas9 off-target effects, we generated a rescue line by stably reintroducing full-length VPS41 with an HA tag (HA-VPS41) using a lentivirus. We confirmed by western blot that the HA-VPS41 expression level matched endogenous levels and by immunofluorescence against the epitope tag that the rescue construct could be detected in every cell (**Figure 1A** and **Supplemental Figure 1B**). As parent INS-1 cells are notoriously heterogeneous, this rescue line represents a good control for our clonal VPS41 KO cells and we have thus used it for all subsequent experiments.

**Figure 1.**
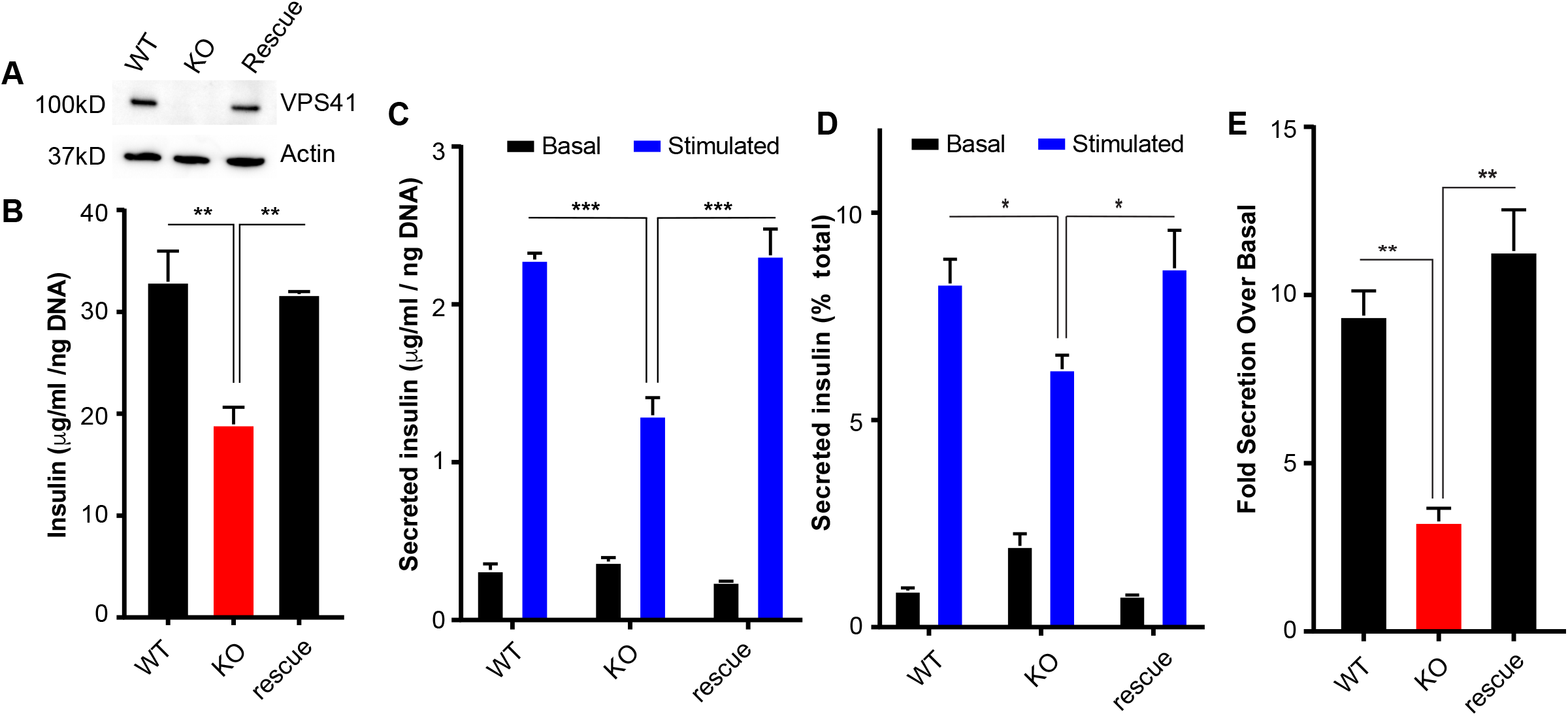
(**A**) Representative western blots showing the levels of VPS41 in WT, VPS41 KO, and HA-VPS41 rescue INS-1 cells. (**B**) Total cellular insulin levels under basal conditions determined by ELISA. (**C**) Basal and stimulated insulin secretion from KO and rescue INS-1 determined by ELISA. (**D**) Insulin secretion relative to cellular insulin stores. (**E**) Fold stimulated insulin secretion over basal insulin secretion. Data indicate mean +/- s.e.m.; n=3 *: p < 0.05, **: p < 0.01, ***: p < 0.001 secretion data analyzed by one-way ANOVA.

We next examined insulin secretion and total cellular insulin content from WT, VPS41 KO and rescue INS-1 cells by ELISA. We observed a reduction in cellular insulin stores as well as in stimulated (17.5mM glucose, 40mM KCl) insulin secretion in the KO cell line (**Figure 1, B and C**). Importantly, after normalizing secreted values to cellular stores, we still observed a defect in regulated insulin secretion in VPS41 KO cells leading to a decrease in the fold stimulation over basal (**Figure 1, D and E**). This indicates that VPS41 not only influences cellular insulin storage, but also influences insulin secretion independently of content.

To test whether the decrease in insulin content is caused by a decrease in insulin production, we measured proinsulin levels in cellular lysates by ELISA. We observed no change between WT and VPS41 KO cells, but VPS41 KO cells displayed slightly increased cellular stores of proinsulin compared to rescue cells, indicating no pronounced decrease in synthesis (**Supplemental Figure 2A**). In addition, VPS41 KO cells showed no difference in proinsulin secretion (**Supplemental Figure 2, B-D**), indicating that proinsulin is not being secreted constitutively. By immunofluorescence, proinsulin distribution remained unchanged between WT, KO and rescue INS-1 cells (**Supplemental Figure 2E**). In sum, we conclude that VPS41 plays a role in cellular insulin content and regulated secretion in β-cells. This is consistent with our previous data in neurons and neuroendocrine cells (Asensio et al., 2013).

### The effect of VPS41 on cellular insulin content and regulated secretion is independent of HOPS

Human patients carrying a point mutation within VPS41 (VPS41^S285P^) exhibit neurological defects without any endocrine dysregulation (van der Welle et al., 2019). As this point-mutant results in impaired HOPS function (van der Welle et al., 2019), we reasoned that this could be a helpful tool to distinguish between HOPS dependent and independent effects of VPS41. Thus, we stably expressed an HA tagged rat VPS41^S284P^ (S285P equivalent in rat) in VPS41 KO INS-1 cells (**Figure 2A)**. We found that these cells displayed a defect in EGF degradation similar to VPS41 KO cells indicating a defect in HOPS (**Figure 2B**) as observed in other cell types (van der Welle et al., 2019). Strikingly, we observed that this mutant rescued insulin secretion and insulin content (**Figure 2, C-F**) phenotypes observed in VPS41 KO INS-1 cells. This finding indicates that the S284P mutation enables VPS41 to function in the regulated secretory pathway independent of the HOPS complex. Consequently, the role of VPS41 in cellular insulin content and regulated secretion is independent of HOPS functionality.

**Figure 2.**
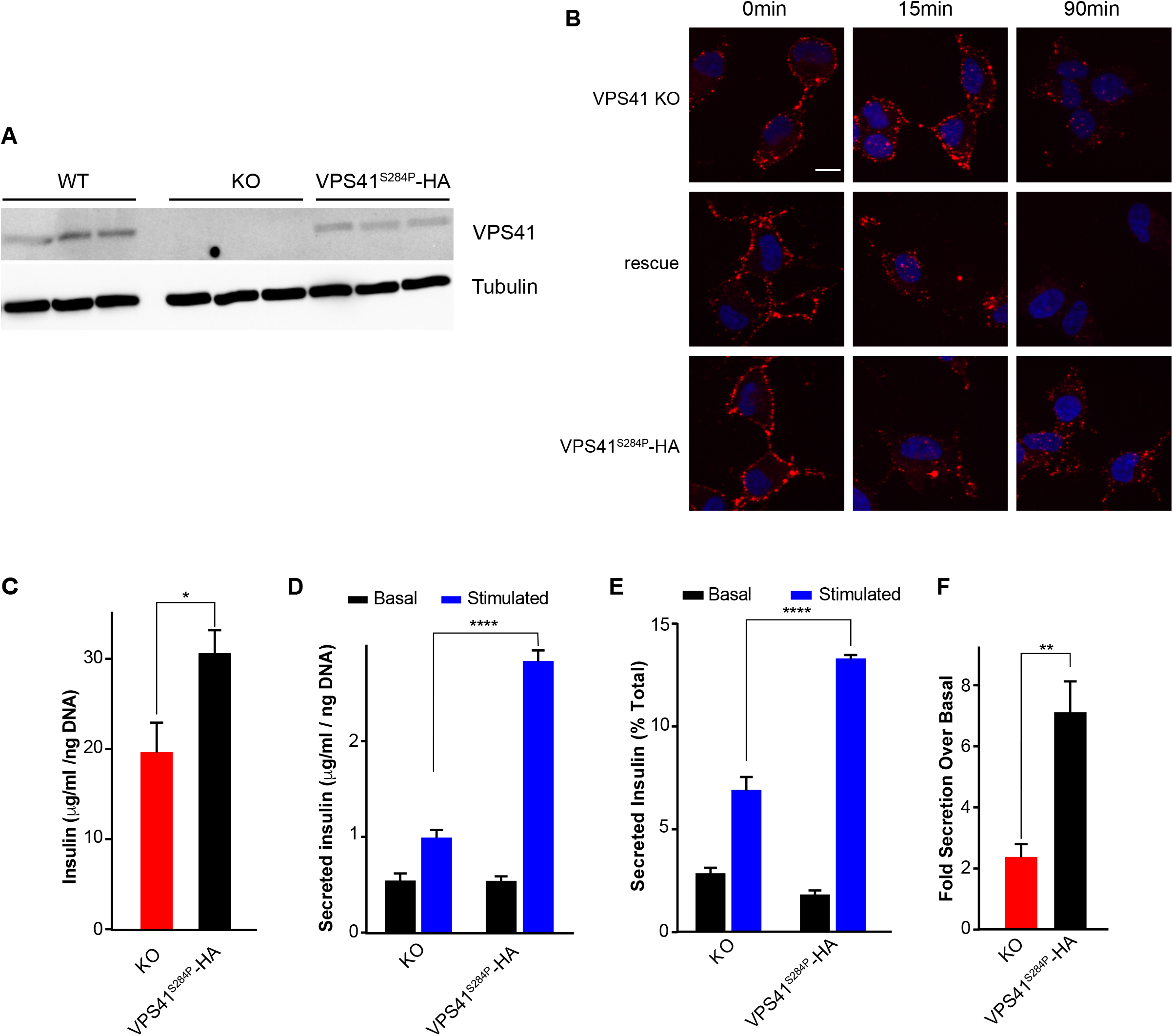
(**A**) Representative western blot showing the levels of VPS41 in WT, VPS41 KO, and HA-VPS41-S284P rescue INS-1 cells. (**B**) Representative images of indicated cells incubated with EGF-Alexa647 before or after a short (15min) or long (90min) chase. (**C**) Total cellular insulin levels of KO and HA-VPS41-S284P rescue INS-1 cells under basal conditions determined by ELISA.(**D**) Basal and stimulated insulin secretion from KO and HA-VPS41-S284P rescue INS-1 cells determined by ELISA. (E) Insulin secretion relative to cellular insulin stores. (F) Fold stimulated insulin secretion over basal insulin secretion. Data indicate mean +/- s.e.m.; n=3 *: p < 0.05, **: p < 0.01, ****: p < 0.001 secretion data analyzed by one-way ANOVA. Scale bar indicates 10μm.

### The absence of VPS41 leads to a decrease in insulin SG number

The decrease in insulin cellular content prompted us to determine whether the absence of VPS41 influences the amount of insulin per vesicle (and thus their density) and/or the number of SGs accumulating intracellularly. For this, we relied first on equilibrium sedimentation through sucrose to separate organelles based on their density and determined insulin distribution within the gradient by ELISA. We observed no change between WT, KO and rescued cells, suggesting no change in SG density (**Figure 3A**). Analysis of thin-sections by EM (**Figure 3, B and C)** or of entire cells by 3D array tomography **(Supplemental Movie 1 and 2)** revealed a decrease in the number of SGs in VPS41 KO cells. Altogether, these data indicate that the absence of VPS41 decreases the number of SGs, but does not influence insulin packaging.

**Figure 3.**
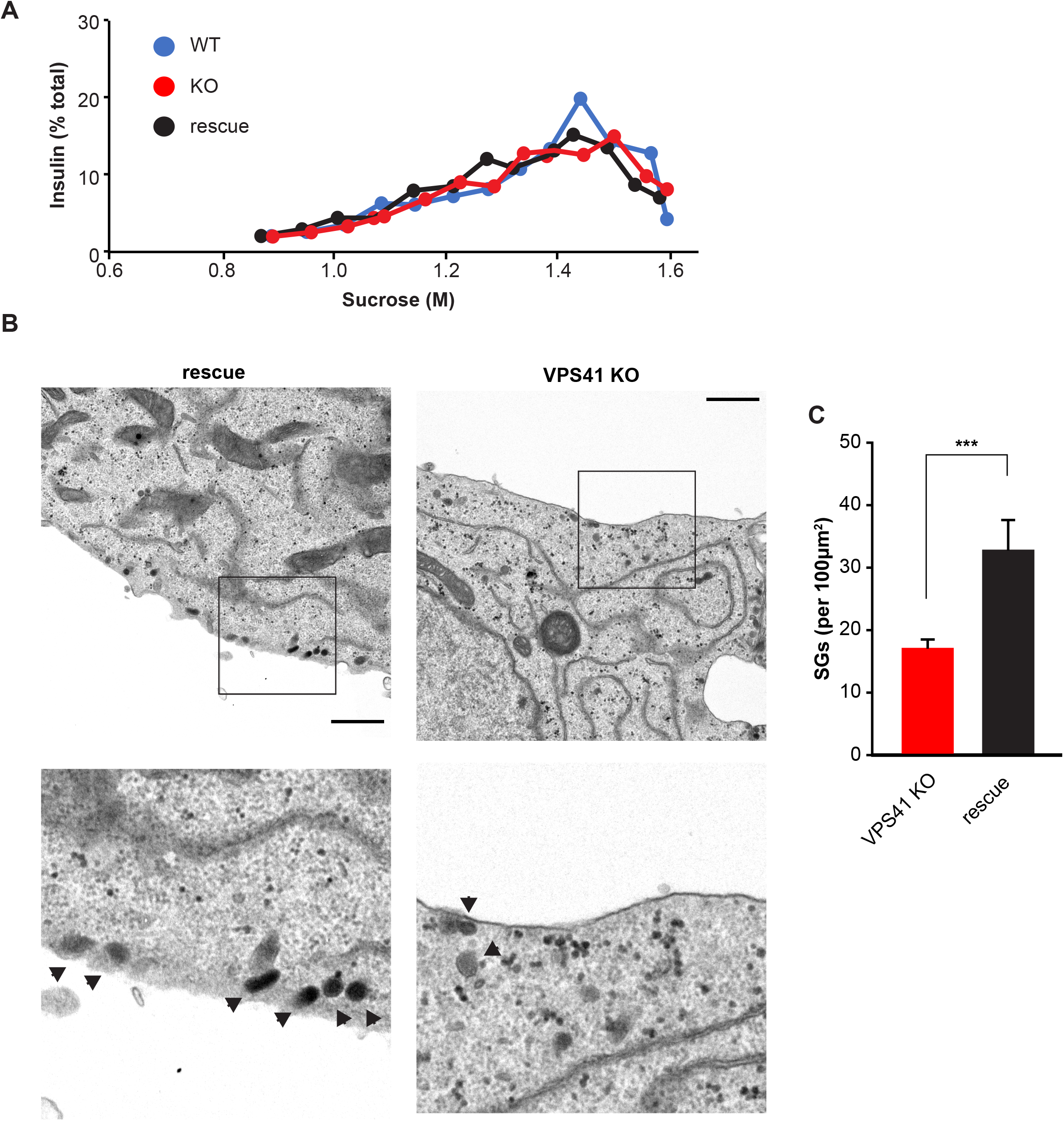
(**A**) Postnuclear supernatants (PNS) obtained from WT, VPS41 KO and HA-VPS41 rescue INS-1 cells were separated by equilibrium sedimentation through 0.6–1.6 M sucrose. Fractions were collected from the top of the gradient. Insulin levels were determined by ELISA in each fraction. The graph indicates the percentage of total gradient insulin from one experiment. Similar results were obtained in an additional independent experiment. (**B**) Representative thin section EM images of VPS41 KO and rescue INS-1 cells. Arrowheads point at SGs. Scale bar indicates 1μm. (**C**) Quantification of total SGs per cell normalized to the area of the cell. Data indicate mean +/- s.e.m.; n= 45 cells for KO and 18 cells for rescue ***: p < 0.001.

### VPS41 KO cells display normal TGN budding kinetics, but transmembrane protein missorting

We next tested whether the decrease in SG number observed in VPS41 KO cells could be a consequence of a budding defect from the TGN. For this, we relied on the ‘retention using selective hook’ (RUSH) system (Boncompain et al., 2012), which allows to synchronize and visualize the movement of cargoes along the secretory pathway. This assay, that we recently optimized for endocrine cells (Hummer et al., 2020), uses localized expression of streptavidin within the endoplasmic reticulum (ER) as a hook to trap a co-expressed secretory cargo fused to streptavidin-binding peptide. Addition of biotin leads to the release of a discrete, synchronized wave of the secretory cargo, in this case, NPY-GFP-SBP, a soluble marker of SGs. Using live-imaging, we monitored the movement of NPY-GFP-SBP along the secretory pathway. Upon addition of biotin, NPY-GFP-SBP trafficked from the ER to the Golgi, followed by its incorporation into post-Golgi carriers in VPS41 KO and rescue cells. By measuring changes in fluorescence, we calculated that the rate of TGN exit remained unchanged between the two cell lines (**Figure 4A**, and **Supplemental Movie 3 and 4**). Next, we determined whether NPY-GFP-SBP sorted to SG properly by determining the degree of colocalization 24 hours after releasing the cargo from its hook. We found that the wave of NPY-GFP-SBP colocalized with endogenous insulin in both cell lines (**Figure 4, B and C**). Furthermore, immunoEM analysis confirmed that proinsulin and insulin sorted properly to SGs (**Supplemental Figure 3**). These data indicate that the budding kinetics and sorting of soluble SG cargoes remain unaffected. We next looked at the behavior of a transmembrane SG marker (phogrin-GFP-SBP) using the RUSH system. Again, we found no difference in TGN exit rates (**Figure 4D)**. However, we observed a striking decrease in phogrin colocalization with endogenous insulin in VPS41 KO cells (**Figure 4, E and F).** This suggests that the absence of VPS41 does not affect TGN budding rates per se but influences SG transmembrane protein composition.

**Figure 4.**
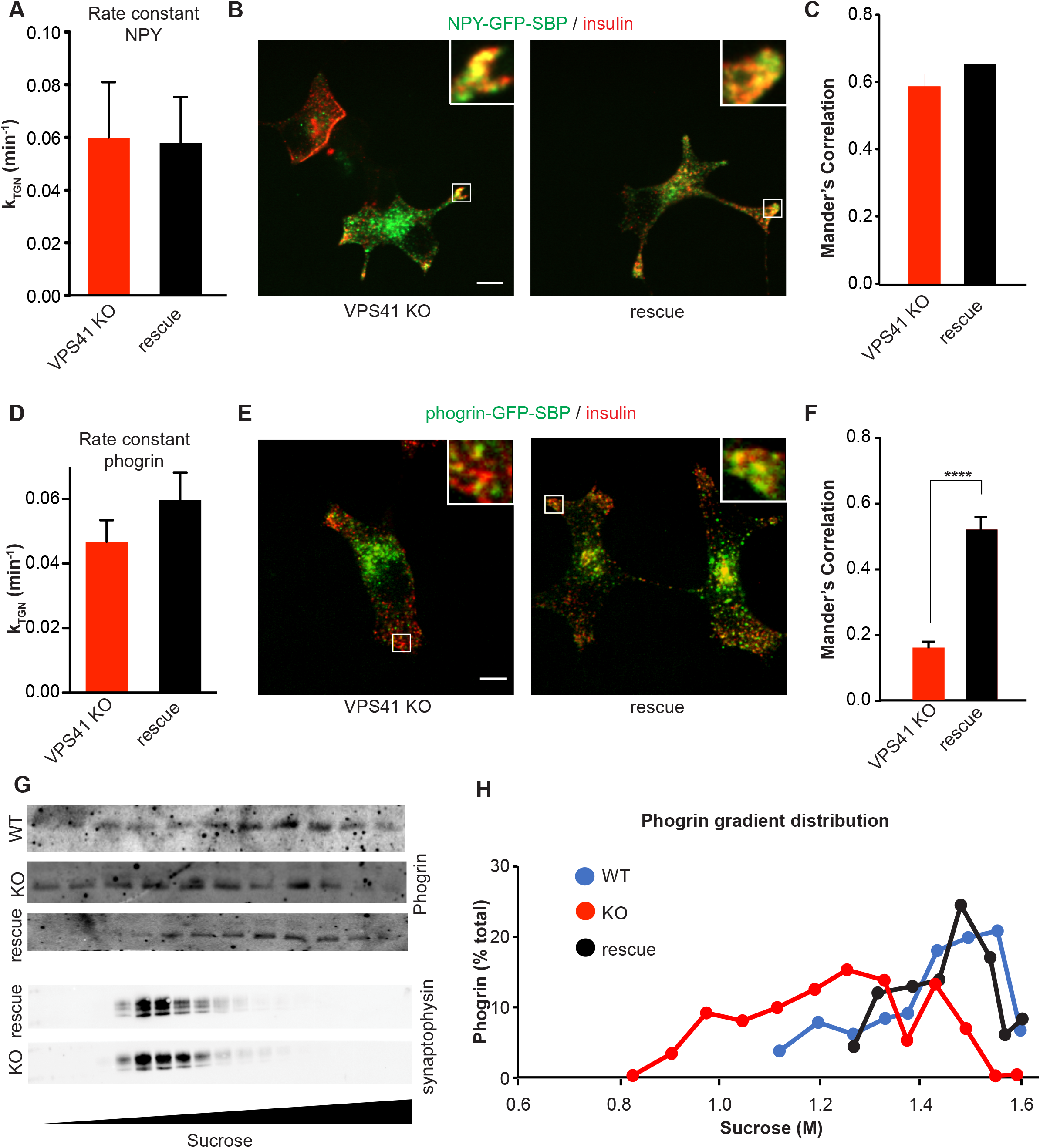
(**A**) TGN rate constants of NPY-GFP-SBP from VPS41 KO and HA-VPS41 rescue INS-1 cells after induction of cargo wave via biotin addition. Data indicate mean +/- s.e.m.; n=9 cells for rescue and n= 5 cells for KO, 2 independent transfections. (**B**) Representative images of NPY-GFP-SBP co-stained with insulin 24h post-biotin addition. Scale bar indicates 10μm (**C**) Manders colocalization coefficient of NPY-GFP-SBP colocalization with insulin signal in VPS41 KO and rescue. Data indicate mean +/- s.e.m.; n=24 cells for rescue and n=24 cells for KO, 2 independent transfections. (**D**) TGN rate constants of phogrin-GFP-SBP from VPS41 KO and HA-VPS41 rescue INS-1 cells after induction of cargo wave via biotin addition. Data indicate mean +/- s.e.m.; n=7 cells for rescue and n= 7 cells for KO, 2 independent transfections. Representative images of Phogrin-GFP-SBP co-stained with insulin 24h post-biotin addition. Scale bar indicates 10μm (**F**) Manders colocalization coefficient of phogrin colocalization with insulin signal in VPS41 KO and rescue. Data indicate mean +/- s.e.m.; n=21 cells for rescue and n= 18 cells for KO, 2 independent transfections. ****: p < 0.0001. (**G**) Postnuclear supernatants (PNS) obtained from WT, VPS41 KO and HA-VPS41 rescue INS-1 cells were separated by equilibrium sedimentation through 0.6–1.6 M sucrose as described in Figure 3. Fractions were assayed for phogrin and synaptophysin (p38) by quantitative fluorescence immunoblotting. (**H**) The graph quantifies phogrin immunoreactivity in each fraction as a percentage of total gradient immunoreactivity from one experiment. Similar results were obtained in an additional independent experiment.

To confirm these findings, we relied on an alternative approach and looked at the distribution of endogenous phogrin after equilibrium sedimentation through sucrose. We observed that phogrin was shifted towards lighter fractions in VPS41 KO cells (**Figure 4, G and H**). This effect appears to be specific to SGs, as sedimentation of synaptophysin (p38), a marker of synaptic-like microvesicles, remained unaffected (**Figure 4G**). These results are consistent with our RUSH data and indicate that phogrin is missorted in absence of VPS41. These data are in line with previous results obtained in PC12 cells (Asensio et al., 2013).

### VPS41 KO cells exhibit defective insulin granule exocytosis

We hypothesized that the absence of VPS41 could lead to the formation of “malfunctioning” SGs that are unable to undergo exocytosis in response to a stimulus, which might explain the defect in regulated insulin secretion. We next tested whether the absence of VPS41 leads to an impairment in SG exocytosis. We transfected VPS41 KO and rescue cells with NPY-pHluorin and monitored exocytosis in real-time using spinning-disk confocal microscopy (**Supplemental Movie 5 and 6**). Under basal conditions, we observed very few exocytosis events across the two cell lines (<1 event/μm^2^/s (x10^-4^)). After stimulation, HA-VPS41 rescue cells showed a robust increase in the number of exocytosis events (55 events/μm^2^/s (x10^-4^)), whereas in the KO cells only 12 events/μm^2^/s (x10^-4^) were observed (**Figure 5A**). Interestingly, we also found that the few exocytosis events observed in the KO cells exhibited very different kinetics. Indeed, in rescue cells, the large majority of events (>80%) lasted less than 1s, but in the KO cells, 50% of events lasted for more than 1s, sometimes up to 9s (**Figure 5B**). Importantly, this exocytosis defect is not caused by an impairment in the excitability of the cells as calcium imaging experiments revealed that VPS41 KO and rescue cells displayed similar responses following depolarization (**Figure 5C**).

**Figure 5.**
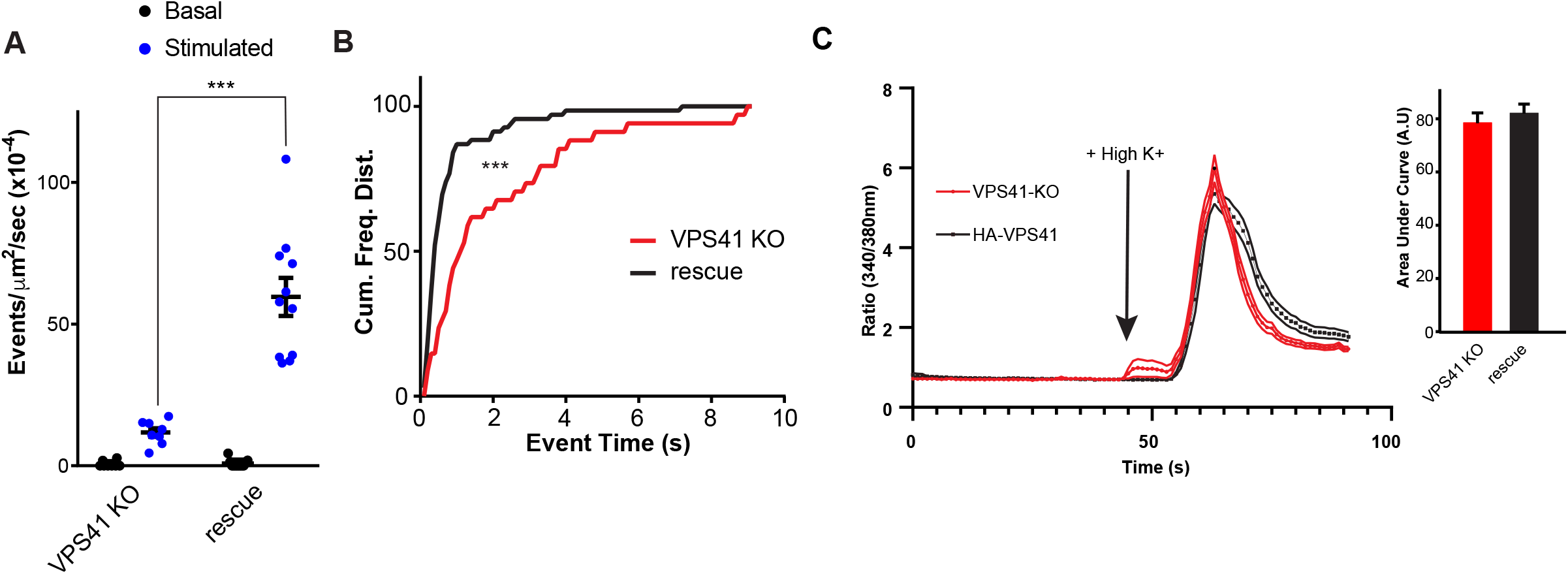
(**A**) Quantification of NPY-pHluorin exocytosis events in VPS41 KO and HA-VPS41 rescue INS-1 cells over a 10s basal or stimulated period. n=3 independent transfections (**B**) Cumulative frequency distribution of NPY-pHluorin exocytosis event duration in VPS41 KO and rescue cells. (n= 34 KO and 69 HA-VPS41 events from 3 independent transfections) (**C**) Average Fura curves in response to depolarization by high K+. Bar graph on the right shows quantification of the area under the curve of calcium imaging (Fura) in response to depolarization (high K^+^) in KO and rescue cells (n= 35 KO and 38 HA-VPS41 cells). Data indicate mean +/- s.e.m.; ***: p < 0.001. Event number analyzed by one-way ANOVA. Event duration analyzed by Kolmogorov Smirnov.

### Mice with β-cell specific deletion of VPS41 develop diabetes due to a defect in insulin secretion and content

To explore the physiological role and significance of VPS41 to glucose homeostasis, we generated β-cell specific VPS41 (βVPS41) KO mice by crossing VPS41 fl/fl animals with mice expressing *Cre* under the control of the insulin promoter; *Ins1-Cre* mice (Thorens et al., 2015). β-cell specificity was confirmed using immunoblotting of different tissues and immunofluorescent staining of islets. Comparison of VPS41 protein levels in lysates prepared from isolated islets revealed a ~75% reduction in βVPS41 KO mice in comparison to their control littermates (**Figure 6, A and B**). The remaining 25% accounts for VPS41 expression in non β-cells of the islets. Importantly, VPS41 levels remained unchanged in whole brain, hypothalamus and liver lysates, indicating that the deletion is specific (**Figure 6, A and B**).

**Figure 6.**
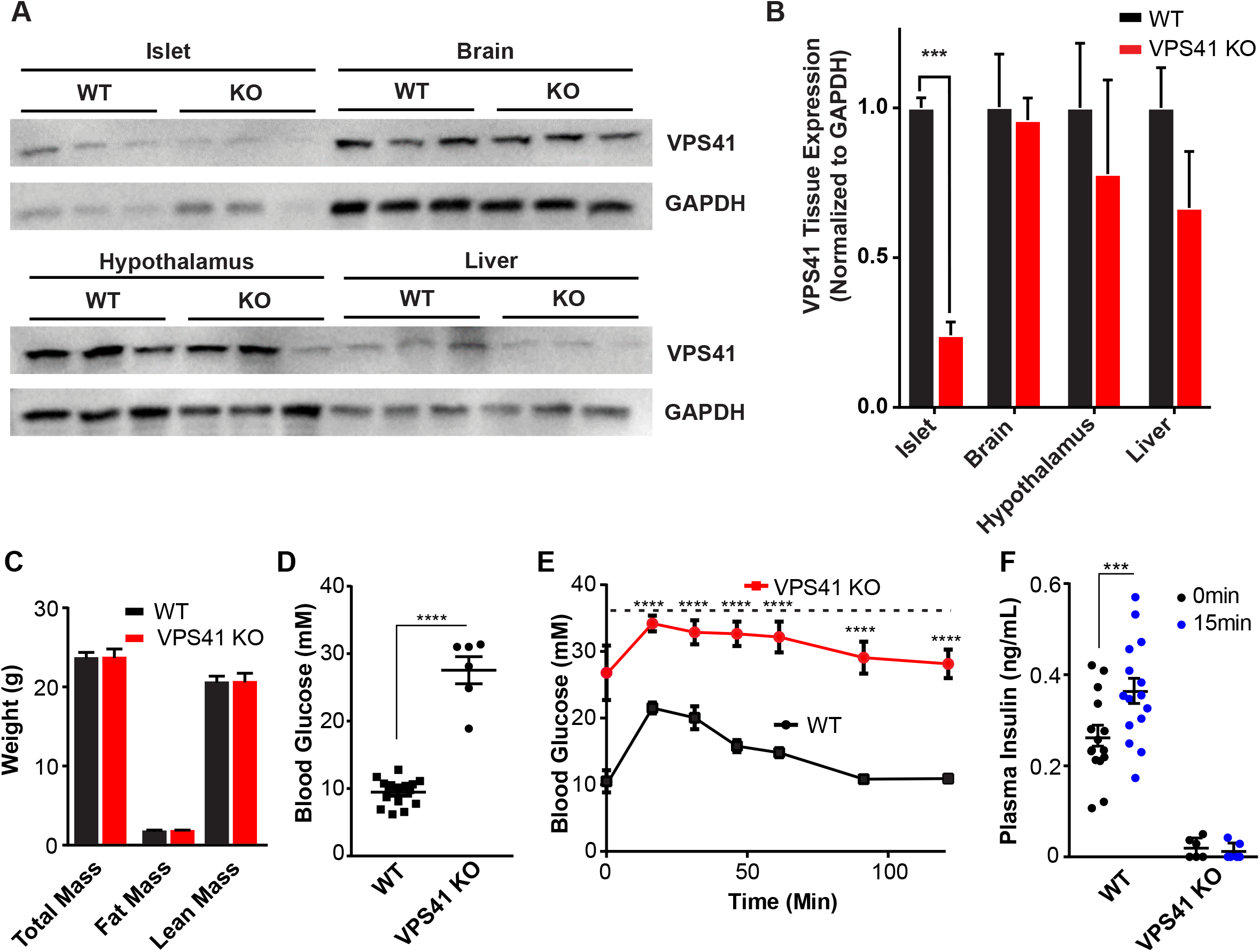
(**A**) Representative western blots of VPS41 protein levels in various tissues from 15-week-old age-matched WT or VPS41 KO mice. (**B**) Quantification of VPS41 protein expression levels normalized to GAPDH. Data indicate mean +/- s.e.m.; n=3 ***: p < 0.001. (**C**) Fat and lean mass measurements of age matched 8-week-old WT and VPS41 KO mice. (**D**) Blood glucose measurement of mice fasted for 8hrs. (E) Blood glucose measurements during glucose tolerance test (GTT). The dotted lines indicate the maximum value of the glucometer. (F) Circulating blood insulin levels before and 15 min after glucose injection. Data indicate mean +/- s.e.m.; KO n=6, WT n=15 ***: p < 0.001, ****: p < 0.0001 secretion data analyzed by one-way ANOVA.

Interestingly, βVPS41 KO mice did not show any changes in body weight, fat or lean mass at 8 (**Figure 6C)** or 15 (**Supplemental Figure 4A**) weeks of age, but these animals develop diabetes characterized by fasting hyperglycemia (**Figure 6D**) and glucose intolerance (**Figure 6E**) without changes in insulin sensitivity (**Supplemental Figure 4E**). At 15 weeks, VPS41 KO mice displayed an even more severe phenotype (**Supplemental Figure 4, B-D**).

The diabetes in βVPS41 KO mice is associated with a pronounced reduction in basal and glucose stimulated insulin levels during the glucose tolerance tests (**Figure 6F**) and severe depletion of insulin in the islets. Immunofluorescence staining of whole pancreas sections revealed a reduction in insulin levels with different degree of severity between cells, but no apparent change in glucagon (**Figure 7A**) and no obvious alteration in the proportion of α- and β-cells. In addition, we isolated islets from 15-week-old animals and determined glucose-stimulated insulin secretion *in vitro*. Consistent with our data in INS-1 cells, we observed a dramatic defect in basal and glucose stimulated insulin secretion from isolated islets (**Figure 7, B and D**), together with a severe reduction in insulin content (**Figure 7C**). Finally, we found a reduction in the number of insulin SGs by EM in islets of VPS41 KO mice (**Figure 7, E and F**).

**Figure 7.**
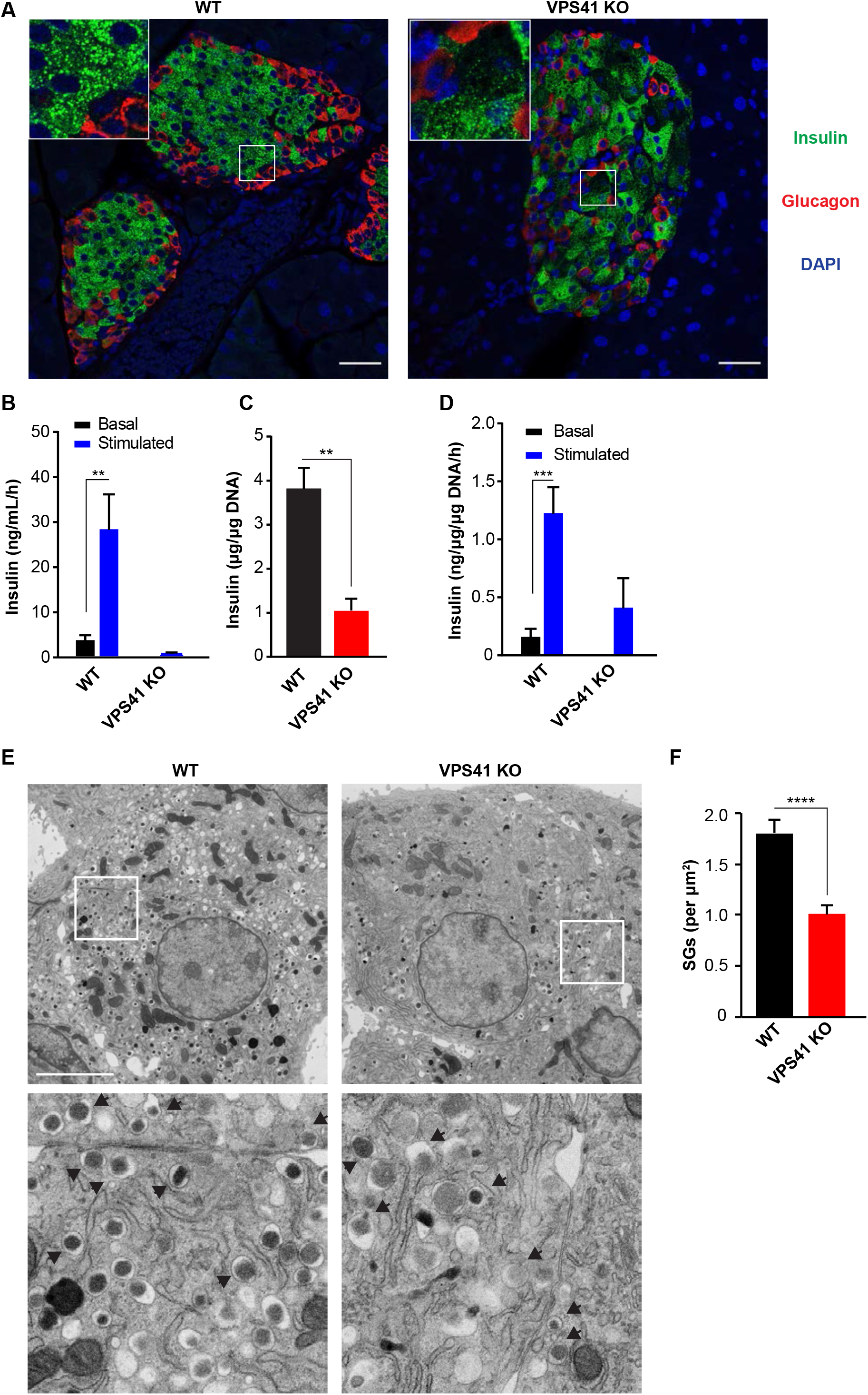
(**A**) Representative immunofluorescence images of whole pancreas slices from WT and VPS41 KO mice stained for insulin (green) and glucagon (red). Scale bar shows 100μm. (**B**) Stimulated insulin secretion from isolated islets from WT and VPS41 KO mice under basal or high glucose conditions. (**C**) Islet insulin content normalized to DNA content, analyzed by t-test. (**D**) Insulin secretion from isolated islets normalized to islet insulin concentration. Data indicate mean +/- s.e.m.; n=3 **: p < 0.01, ***: p < 0.001, secretion data analyzed by one-way ANOVA. (E) Representative thin section EM images of islets from WT and VPS41 KO mice. Arrowheads point at SGs. Scale bar indicates 5μm. (F) Quantification of total SGs per cell normalized to the area of the cell. Data indicate mean +/- s.e.m.; n= 128 cells for WT from 3 animals and 141 cells from 3 animals ****: p < 0.0001.

In conclusion, we find that deletion of VPS41 specifically in β-cells leads to an overtly diabetic mouse, due to decreased insulin secretion and content. However, this mouse does not exhibit obvious defects in glucagon levels or islet architecture, suggesting again that VPS41 is playing a specific role in the regulation of regulated insulin storage and release, both *in vitro* and *in vivo*.

## Discussion

Loss of VPS41 from neuroendocrine PC12 cells and neurons leads to defects in peptidergic regulated secretion (Asensio et al., 2013). SGs can still form in these cells, but they show altered size and morphology. We now show that absence of VPS41 also leads to defects in insulin regulated secretion and cellular content in β-cells. Thus, although SGs display a wide range of size and cargo content across specialized secretory cells, some of the molecular mechanisms controlling their formation seem conserved between neurons, neuroendocrine and endocrine cells.

We also find that the S285P human mutation, which is unable to rescue HOPS function (van der Welle et al., 2019), restores insulin secretion completely, indicating that it is actually possible to isolate HOPS dependent and independent functions of VPS41. Furthermore, this observation also helps explain why patients bearing this mutation have no observable endocrine associated defects. It might also show that the HOPS independent function of VPS41 is necessary for survival, since VPS41 KO mice die early during embryogenesis (Aoyama et al., 2012). In summary, our data indicate that VPS41 is required for insulin secretion and intracellular storage, and this effect is independent of HOPS.

As the NPY-pHluorin assay did not reveal any changes in the number of basal exocytosis events and as insulin basal secretion remained unchanged in absence of VPS41, it is unlikely that rerouting of soluble regulated cargoes to the constitutive secretory pathway accounts for the decrease in SGs. We also did not observe a decrease in proinsulin cellular levels, suggesting that insulin production remains unaffected. Finally, the results from the RUSH experiments indicate that the lack of VPS41 does not impair cargo budding. This finding is further substantiated by the lack of excessive accumulation of proinsulin at the Golgi in VPS41 KO cells compared to WT and rescue cells by immunofluorescence. It is thus tempting to speculate that the lack of VPS41 leads to degradation of SGs. Indeed, degradation is an important mechanism by which β-cells maintain insulin granule homeostasis (Brereton et al., 2014; Marsh et al., 2007). Several modes of SG degradation have been described over the years, including macroautophagy, crinophagy and stress-induced granule degradation (SINGD), which both involves the direct fusion of granules with lysosomes, and Golgi membrane-associated degradation (GOMED) (Halban, 1991; Orci et al., 1984; Pasquier et al., 2019; Riahi et al., 2016; Yamaguchi et al., 2016). As crinophagy and macroautophagy are HOPS-dependent processes (Csizmadia et al., 2018; Jiang et al., 2014), these pathways probably do not contribute much to granule turnover as they are likely to be defective in VPS41 KO cells. In the future, it will be interesting to test whether there is an increase in SINGD and/or GOMED in VPS41 KO cells once the molecular mechanisms for these processes are better understood.

In addition, independently from the decrease in cellular insulin content, our results show that the absence of VPS41 leads to a defect in regulated secretion of insulin. Our data suggest that VPS41 might control sorting of transmembrane proteins, which ultimately could influence SG release properties. We have previously proposed that VPS41 functions at the TGN during budding of SGs in neuroendocrine cells (Asensio et al., 2013). Consistent with this, a pool of VPS41 localizes to the TGN (Asensio et al., 2013; Pols et al., 2013). Alternatively, VPS41 might contribute to SG biogenesis by influencing maturation after budding. Indeed, several cytosolic factors that have recently been implicated in SG biogenesis are typically associated with endocytic and retrograde pathways (Asensio et al., 2010; Cao et al., 2013; Holst et al., 2013; Paquin et al., 2016; Pinheiro et al., 2014; Sasidharan et al., 2012; Sirkis et al., 2013; Zhang et al., 2017). The EARP complex and its interacting partner EIPR-1 for example, contribute to SG cargo sorting (Topalidou et al., 2020; Topalidou et al., 2016). It is thus tempting to speculate that retrograde pathways might contribute to SG maturation and that VPS41 could influence this process. Interestingly, in addition to a reduction in the number of exocytotic events after stimulation, we observed a change in the release properties of SGs in VPS41 KO cells with an increase in the duration of the events. Thus, we can’t exclude that VPS41 contributes to SG exocytosis directly by interacting with SNAREs or other factors of the exocytosis machinery.

Islets of BTBR mice, which are prone to develop diabetes, exhibit decreased levels of VPS41 gene expression (Keller et al., 2008). In agreement with this, we now show that mice with a specific deletion of VPS41 in β-cells are overtly diabetic. Indeed, these animals display extreme fasting hyperglycemia and are glucose intolerant due to a defect in insulin content and secretion. Disruption of lysosomal genes such as raptor leads to a very similar phenotype, however, in this case, the defect in secretion seems to be linked to β-cell maturation and early onset decreases in β-cell mass (Ni et al., 2017). In contrast, we observed no obvious difference in islet size and morphology in absence of VPS41. Our data indicate that VPS41 plays a significant role for glucose homeostasis and normal physiology. Our findings, together with the identification of several point-mutations associated with type 2 diabetes in humans (Type_2_Diabetes_Knowledge_Portal), further underline the importance of membrane trafficking in health and disease (Gilleron et al., 2019). In particular, our data emphasize the physiological significance of understanding the mechanisms of SG formation at the molecular level. Unraveling the role of cytosolic factors in this process could help reveal potential therapeutic targets for the treatment of metabolic and endocrine associated diseases.

## Methods

### Molecular Biology and Plasmids

The human codon-optimized Cas9 and chimeric guide RNA expression plasmid (pX459v2) developed by the Zhang lab were obtained from Addgene (Cong et al., 2013). To generate gRNA plasmids, a pair of annealed oligos (20 base pairs) were ligated into the single guide RNA scaffold of pX459v2. The following gRNAs sequences were used: Forward (rat): 5’-CACCGACTCTCAGACTGAGCTATGG-3’; Reverse (rat): 5’-AAACCCATAGCTCAGTCTGAGAGTC-3’ to generate the INS1 VPS41 KO line. Rat VPS41 lentiviral plasmid was generated by amplifying VPS41 from rat cDNA using the following primers: WT Forward: 5’-CCTCCATAGAAGACACCGACTCTAGACACCATGGCGGAAGCAGAGGAG-3’; WT Reverse: 5’-TATGGGTAACCCCCAGATCCACCGGTCTTCTTCATCTCCAGGATGGCA-3’.

The PCR products were then subcloned by Gibson ligation into pLenti-CMV-GFP-Puro. pLenti-CMV-GFP-Puro (Witwicka et al., 2015) was a gift from Paul Odgren (Addgene plasmid #73582). The following primers were used to genotype VPS41 KO INS-1 cells: rVPS41-F: 5’-ATATGGTACCTGCAGAGAGGACTAAACGAGA-3’; rVPS41-R 5’-ATATGAGCTCATGGACGTTGCCATCCAAGT-3’. To test for the presence of indels, the resulting PCR products were ligated into pBlueScript II KS. Isolated plasmids from 10 random colonies were then analyzed for the presence of indels by Sanger sequencing. ON-TARGETplus siRNA pools were obtained from Dharmacon (VPS41: 306991) and used as previously described (Asensio et al., 2013). All RUSH constructs used were generated using pEGFPC3 as a backbone. STR-Linker-KDEL was used as the ER hook. NPY-eGFP-SBP, NPY-pHluorin-SBP, phogrin-GFP-SBP, and VMAT2-GFP-SBP were generated and used as cargo plasmids (Hummer et al., 2020).

### Cell Culture and Lentiviral Production

INS-1 cells were maintained in RPMI supplemented with 10% FBS, 1mM Na-Pyruvate, 50uM β-mercaptoethanol (β–ME) under 5% CO_2_at 37°C. INS1 were transfected using Lipofectamine 2000 or Fugene-HD as per manufacturer’s instructions. HEK293T cells were maintained in DMEM supplemented with 10% FBS under 5% CO_2_at 37°C. Lentivirus was produced by transfecting HEK293T cells with pLenti-CMV-Puro, psPAX2, and pVSVG using 1mg/mL PEI. CRISPR-Cas9 KO lines were generated by transfection of WT INS-1 cells with pX459v2 containing the guides of interest with Fugene-HD as per manufacturer’s instructions. 2 days post-transfection, cells were treated with 4ug/ml puromycin for 2 days. Following selection, KO clones were isolated as single clones and expanded prior to sequencing of indels. Cells were routinely tested for the absence of mycoplasma.

### Insulin Secretion Assays

INS-1 insulin secretion was monitored by ELISA (ThermoFisher). Cells were plated at 200k cells per 24 well. Prior to stimulation, cells were moved to complete media containing 5mM glucose 18hrs prior, and to KRB buffer supplemented with 1.5mM glucose 2hrs prior and maintained with 5% CO_2_ at 37°C. Cells were either stimulated in 250ul KRB supplemented with 40mM KCl and 16.7mM glucose or in low glucose KRB for 2hrs. The secreted fraction was removed and spun @1k xg for 5min to remove cellular debris. Cells were lysed in 500ul lysis buffer (50mM TRIS-HCl, 300mM NaCl, 2% v/v TritonX-100) supplemented with 1mM PMSF and 1X protease inhibitor cocktail on ice for 5min. Cellular debris removed by spinning at 21k xg for 10min at 4°C.

ELISA was performed as per manufacturer instructions with the secreted fraction being diluted 1:2 and the cellular fraction being diluted 1:10. Proinsulin secretion was monitored by ELISA (Mercodia). Samples, both cellular and secreted, were diluted 1:2 for the assay. Secretion was normalized to cell content by measuring DNA in the cellular fraction. Blood insulin and islet insulin secretion ELISA was conducted according to manufacturer’s protocol (Crystal Chem Ultra Sensitive Mouse ELISA)

### Thin-section Transmission Electron Microscopy and Array Tomography

Coverslips were rinsed and fixed in 2.5% glutaraldehyde and 2% paraformaldehyde in 0.15M cacodylate buffer pH 7.4 with 2 mM calcium chloride warmed to 37°C for 5 minutes in the incubator at 37°C. After this, the samples were transferred to a refrigerator at 4°C in the same fixative solution. Once ready to be processed, the coverslips were rinsed in 0.15 M cacodylate buffer 3 times for 10 minutes each and subjected to a secondary fixation for one hour in 1% osmium tetroxide / 0.3% potassium ferrocyanide in cacodylate buffer on ice. Following this, samples were then washed in ultrapure water 3 times for 10 minutes each and *en bloc* stained for 1 hour with 2% aqueous uranyl acetate. After staining was complete, samples were briefly washed in ultrapure water, dehydrated in a graded acetone series (50%, 70%, 90%, 100% x2) for 10 minutes in each step, infiltrated with microwave assistance (Pelco BioWave Pro, Redding, CA) into LX112 resin, and flat embedded between two slides that had been coated with PTFE release agent (Miller-Stephenson #MS-143XD, Danbury, CT). Samples were cured in an oven at 60°C for 48 hours. Once resin was cured, slides were separated and the aclar coverslips peeled off. A small region was then cut out and glued onto a blank stub with epoxy. For TEM imaging, 70nm thin sections were then cut using a diamond knife with a 35-degree angle (Diatome, Hatfield, PA), placed onto grids and imaged on a JEOL 1400-Plus TEM at 120kV. For Array Tomography, approximately 90, 80nm sections were taken in serial and placed onto a silicon wafer and were imaged on a FE-SEM (Zeiss Merlin, Oberkochen, Germany) using the BSD detector. The SEM was operated at 8 KeV and a probe current of 0.9 nA. The same 122 μm field of view in each slice was imaged using the Array Tomography routine in ATLAS 5 (FIBICS, Ottawa, Ca) at 10nm/pixel resolution at 5ns dwell time with 4 line averaging.

### Immunofluorescence and Spinning Disk Confocal Microscopy

INS-1 cells were rinsed with PBS and fixed in 4% paraformaldehyde in PBS and incubated for 20 min at room temperature. Cells were permeabilized in PBS containing 0.1% Triton X-100 for 10 min at room temperature and blocked in PBS containing 2% BSA, 1% fish skin gelatin, and 0.02% saponin. Primary and secondary antibodies were diluted as reported in Table 2. Images were acquired using a custom-built Nikon spinning disk at a resolution of 512 × 512 pixels with a 63 × objective (Oil Plan Apo NA 1.49) and an ImageEM X2 EM-CCD camera (Hamamatsu, Japan). Antibodies used are listed in **Table 1**.

**Table 1.** Antibodies used in this study

| Antibody Target | Source | Dilution (Application) |
| --- | --- | --- |
| VPS41 (mouse) | Santa Cruz Biotech. (D-12) | 1:100 (Western) |
| Tubulin (mouse) | DSHB (E-7) | 1:500 (Western) |
| Actin (mouse) | DSHB (JLA20) | 1:500 (Western) |
| Synaptophysin (mouse) | Synaptic Systems (MMS-618R) | 1:2000 (Western) |
| PC1/3 (rabbit) | Richard Mains (Gift) | 1:500 (Western) |
| TFE3 (rabbit) | Cell Signaling Tech. (14779) | 1:500 (IF) |
| Insulin (mouse) | Sigma Aldrich (K36AC10) | 1:1000 (IF) |
| HA (rat) | Sigma Aldrich (3F10) | 1:1000 (IF) |
| Phogrin (rabbit) | Zippy from John Hutton | 1:100 (western) |
| Proinsulin (mouse) | DHSB (GS-9A8) | 1:500 (IF) |

### NPY-pHluorin Exocytosis Imaging

INS-1 cells were co-transfected with NPY-pHluorin and NPY-mCherry. At 1 day after transfection, cells were transferred to poly-l-lysine coated 22 mm glass coverslips. After an additional 2 days, cells were moved to KRB buffer supplemented with 1.5mM glucose 2hr prior to imaging and maintained with 5% CO_2_at 37°C. After reset, cells were washed with KRB buffer and coverslips were transferred to an open imaging chamber (Life Technologies). Cells were imaged close to the coverslips, focusing on the plasma membrane (determined by the presence of NPY-mCherry positive plasma membrane docked-vesicles), using a custom-built Nikon spinning disk at a resolution of 512 × 512 pixels. Images were collected for 100 ms at 10 Hz at room temperature with a 63 × objective (Oil Plan Apo NA 1.49) and an ImageEM X2 EM-CCD camera (Hamamatsu, Japan). Following baseline data collection (15 s), an equal volume of 2x KRB buffer containing 114 mM KCl (60mM final) and 32.65mM Glucose (16.7mM final) was added to stimulate secretion and cells were imaged for an additional 45 s. At the end of the experiment, cells were incubated with KRB solution containing 50 mM NH4Cl, pH 7.4, to reveal total fluorescence and to confirm that the imaged cells were indeed transfected. Movies were acquired in MicroManager (UCSF) and exported as tif files. Events were quantified within the first 10s of basal stimulation and between 10s post stimulation (15s after high K+ addition). Event length was calculated based on the time the event remained at or above 20% above baseline signal. For Fura imaging, cells were plated onto poly-l-lysine (Sigma) coated imaging dishes and allowed to adhere in media for 2 days. To load FURA-2, AM (Life Technologies), a basal KRB solution with 500nM FURA was added to the cells. Cells were incubated in the dark at room temperature for 30 minutes, the FURA-2, AM solution was removed, and cells washed briefly with basal solution was added to the dish. Cells were imaged using a CoolSNAP HQ2 camera with 6.21 μm/pixel using a 40x oil objective (1.3 N.A.). Images were binned (2×2) and cells were alternatively excited by 340 nm and 380 nm light every 500 ms with a 100 ms interval between exposures. Cells were imaged for 45s under basal conditions prior to perfusing cells with a 90mM K+ KRB solution for 45s.

### RUSH Assays

Constructs used for this study have been described before (Hummer et al., 2020). For RUSH, a ratio of 3:1 hook to cargo ratio was determined to be ideal. Lipofectamine 2000 was used to transfect cells. Experiments were performed 36-48h after transfection. For live imaging, cells were transfected in 24-well dishes, 24h post transfection cells were trypsinized and re-plated onto a 35mm glass bottom imaging dish (live imaging) or 10mm glass coverslips coated with PLL. Cells were imaged 24h after being re-plated. For RUSH, INS-1 cells were grown in biotin and phenol-red free RPMI media supplemented with 2.4mM sodium bicarbonate and 25mM HEPES. 50μM biotin addition was used to initiate cargo release. Live RUSH movies were imaged on an Evos FL Auto 2 in an environmentally controlled imaging chamber (37C, 5% CO_2_, 20% humidity). Cells were imaged every 2min.

### Density Gradient Fractionation

Equilibrium sedimentation through sucrose was performed as previously described (Asensio et al., 2013). Briefly, a postnuclear supernatant was prepared from INS-1 cells by homogenization with a ball bearing device (12 μm clearance), loaded onto a 0.6–1.6 M continuous sucrose gradient, and sedimented at 30,000 rpm in an SW41 rotor for 14–16 h at 4°C. Fractions (750μl each) were collected from the top and were analyzed by quantitative fluorescence immunoblotting using a FluoChem R imager (ProteinSimple). Insulin concentration in the fractions was determined by ELISA as before, after diluting fraction 1:4 in ELISA lysis buffer for 5min, then diluting sample 1:5 in ELISA diluent.

### EGF Degradation Assay

INS-1 cells plated on PLL coated glass coverslips, were washed twice with PBS and starved of serum for 2 h in DMEM with 0.1% BSA (GoldBio). Following starvation, cells were washed twice with ice-cold PBS on ice and incubated with EGF conjugated with Alexa-Fluor 647 (ThermoFisher) at a final concentration of 1 μg/ml for 1 h on ice taking precautions to protect from light. Excess unbound EGF was removed by washing with ice-cold PBS in presence of 0.5% BSA.

Cells were chased for the indicated times at 37C before fixation with 4% PFA in PBS for 30 min on ice. Cells were imaged using a spinning disk confocal microscope.

### Immuno-electron microscopy

VPS41 KO or HA-VPS41 expressing INS-cells were grown in a T25 flask and fixed using freshly made 4% Formaldehyde (FA), 0.4% Glutaraldehyde (GA) in 0.1M phosphate buffer (pH 7.4) by adding an equal amount of fixative to the medium for 5 min. Cells were postfixed using 2% FA, 0.2% GA in 0.1M phosphate buffer for 2 hours and stored in 1% FA at 4°C. Ultrathin cryosectioning and immunogold labelling were performed as previously described (Slot & Geuze, 2007). Proinsulin and insulin were detected using mouse anti-Proinsulin (GS9A8, Madsen, 1:10000) and rabbit anti-Insulin (Kuliawat et al., 1997) (kindly provided by Dr. P. Arvan, 1:10000) respectively. Primary and bridging (rabbit anti-mouse, 610-4120 Rockland, 1:250) antibodies were detected by protein A–10 nm gold particles (Cell Microscopy Center, Utrecht, The Netherlands).

### Mouse Housing and Breeding

Mice are bred at LAS Facility, Bosch Institute, University of Sydney, Australia and transferred to the Charles Perkins Centre, University of Sydney for experimental procedures. All mice are fed a standard laboratory chow (13% calories from fat, 65% carbohydrate, 22% protein from Gordon’s Specialty Stock Feeds, Yanderra, New South Wales, Australia).

### Mouse Breeding

VPS41^fl/fl^ mice (Aoyama et al., 2012) were crossed with Ins1^cre^ mice (Jax strain 026801) to generate third generation VPS41^fl/fl^, Ins1^cre/+^ mice and their littermate controls for experiments.

Mice were bred at Bosch Animal Facility, University of Sydney, Australia and transferred to the Animal Facilities at the Charles Perkins Centre for experimental use. Mice were maintained on a 12-h light/dark cycle (0700/1900 h) and given ad libitum access to food and water. Experiments were carried out in accordance with the National Health and Medical Research Council (Australia) guidelines for animal research and were approved by the University of Sydney Animal Experimentation Ethics Committees.

### Glucose Tolerance Test

Mice are fasted 5 hours, then analyzed prior to IP-GTT by EchoMRI to calculate dose of 2mg/kg glucose to lean mass. Mice are tail tipped for basal blood glucose and insulin measurement, then injected with a 25% glucose solution in injectable water via intraperitoneal route. Blood is collected at 15 minutes for fed insulin measurement, and at the following timepoints: 15, 30, 45, 60, 90, 120 minutes for glucose measurements.

### Static Islet Insulin Secretion

Following islet isolation, islets recovered in recovery medium (RPMI, 10% FBS, 11.1mM glucose) for 1hr at 37°C. 10 islets per well (in triplicate) were placed into KRB with no glucose. Islets were transferred to KRB solution containing 2.8mM glucose, as basal incubation, or 16.7mM glucose KRB, as stimulatory and incubate at 37°C for 1 hr. Islets were subsequently lysed into Islet Lysis Buffer (100 mM Tris, 300 mM NaCl, 10 mM NaF, 2 mM sodium orthovanadate). Secreted and cellular insulin content was measured via ELISA (Crystal Chem), while total DNA content was measured by PicoGreen DNA Assay (Quant-iT Picogreen DNA kit, Thermo Fisher Scientific).

### Fluorescent Immunohistochemistry

Whole pancreas was imbedded with paraffin, prior to thin sectioning. Thin sections of pancreas are deparaffinized and rehydrated for 3min in the following solutions: 2x xylene rinse solution, 1x 50% xylene / 50% ethanol, 1x 100% ethanol, 1x 95% ethanol, 1x 80% ethanol, 1x 75% ethanol, 1x 50% ethanol, 1x ddH_2_O. Slides were washed 3min 2x in Wash Buffer (PBS with 0.1% BSA, 0.01% NaN_3_), Non-specific antibody association was blocked with 2 drops of blocking solution (DAKO). Primary antibody was diluted in dilution buffer (DAKO), 100 μl per section, and incubated overnight @4C in a humidified chamber. Slices were washed 3x with Wash Buffer for 3min each. Secondary antibody was diluted in dilution buffer, 100 μl per slice was added @RT for 1hr. Slices were rinsed with wash buffer 2x for 3min and with PBS 2x for 3min. Slice were dried for 5min @RT, before mounting with Prolong Diamond Antifade Mountant (Invitrogen).

### Statistics

Unless indicated otherwise, all statistical analysis was performed using the two-tailed Student’s t test. Statistical analyses were conducted using Excel or Prism.

## Figure preparation

Images were processed using ImageJ; any changes in brightness and contrast were identical between samples meant for comparison.

## Author contributions

CHB conducted in vitro experiments, acquired and analyzed data, and wrote the manuscript. BY and YSA conducted in vivo experiments, acquired and analyzed data. AR, DM, JT, RW, TV conducted in vitro experiments, acquired and analyzed data. DC identified the S285P human mutation. GWS and RGK conducted the thin section EM, MRF and POB conducted the array tomography experiments. JAJF, MAK and JK supervised the study and edited the manuscript. CSA designed the study, analyzed data, supervised the study and wrote the manuscript.

## Acknowledgments

We thank Peter Arvan for the INS-1 cell line and Yoh Wada for the VPS41^fl/fl^ mice; PC1/3 antibody was a gift from Dick Mains and Betty Eipper. The Washington University Center for Cellular Imaging (WUCCI) gratefully acknowledges support from Washington University School of Medicine, The Children’s Discovery Institute of Washington University and St. Louis Children’s Hospital (CDI-CORE-2015-505 and CDI-CORE-2019-813 to JAJF), the Foundation for Barnes-Jewish Hospital (3770 to JAJF) and the Washington University Diabetes Research Center (DRC) (P30 DK020579). MAK is supported by a Jennie Mackenzie Philanthropic Fellowship, University of Sydney. Y.A. is a recipient of the Australian Postgraduate Scholarship. JK and RvdW are supported by the Deutsche Forschungs Gemeinshaft (DFG) FOR2625. This work was supported by American Diabetes Association grant #1-17-JDF-064 and by NIH grants R01 GM124035 and R15 GM116096 to CSA.

## Notes

The authors have declared that no conflict of interest exists.

### Competing Interest Statement

The authors have declared no competing interest.

## References

Ahras, M., Otto, G. P., & Tooze, S. A. (2006). Synaptotagmin IV is necessary for the maturation of secretory granules in PC12 cells. J Cell Biol, 173(2), 241–251. doi:10.1083/jcb.200506163

Aoyama, M., Sun-Wada, G. H., Yamamoto, A., Yamamoto, M., Hamada, H., & Wada, Y. (2012). Spatial restriction of bone morphogenetic protein signaling in mouse gastrula through the mVam2-dependent endocytic pathway. Dev Cell, 22(6), 1163–1175. doi:10.1016/j.devcel.2012.05.009

Asensio, C. S., Sirkis, D. W., & Edwards, R. H. (2010). RNAi screen identifies a role for adaptor protein AP-3 in sorting to the regulated secretory pathway. J Cell Biol, 191(6), 1173–1187. doi:jcb.201006131 [pii] 10.1083/jcb.201006131

Asensio, C. S., Sirkis, D. W., Maas, J. W., Jr., Egami, K., To, T. L., Brodsky, F. M., . . . Edwards, R. H. (2013). Self-assembly of VPS41 promotes sorting required for biogenesis of the regulated secretory pathway. Dev Cell, 27(4), 425–437. doi:10.1016/j.devcel.2013.10.007 S1534-5807(13)00604-7 [pii]

Balderhaar, H. J., & Ungermann, C. (2013). CORVET and HOPS tethering complexes - coordinators of endosome and lysosome fusion. J Cell Sci, 126(Pt 6), 1307–1316. doi:10.1242/jcs.107805

Boncompain, G., Divoux, S., Gareil, N., de Forges, H., Lescure, A., Latreche, L., . . . Perez, F. (2012). Synchronization of secretory protein traffic in populations of cells. Nat Methods, 9(5), 493–498. doi:10.1038/nmeth.1928

Borgonovo, B., Ouwendijk, J., & Solimena, M. (2006). Biogenesis of secretory granules. Curr Opin Cell Biol, 18(4), 365–370. doi:10.1016/j.ceb.2006.06.010

Brereton, M. F., Iberl, M., Shimomura, K., Zhang, Q., Adriaenssens, A. E., Proks, P., … Ashcroft, F. M. (2014). Reversible changes in pancreatic islet structure and function produced by elevated blood glucose. Nat Commun, 5, 4639. doi:10.1038/ncomms5639

Cao, M., Mao, Z., Kam, C., Xiao, N., Cao, X., Shen, C., … Xia, J. (2013). PICK1 and ICA69 control insulin granule trafficking and their deficiencies lead to impaired glucose tolerance. PLoSBiol, 11(4), e1001541. doi:10.1371/journal.pbio.1001541 PBIOLOGY-D-12-01903 [pii]

Cong, L., Ran, F. A., Cox, D., Lin, S., Barretto, R., Habib, N., … Zhang, F. (2013). Multiplex genome engineering using CRISPR/Cas systems. Science, 339(6121), 819–823. doi:10.1126/science.1231143 science.1231143 [pii]

Csizmadia, T., Lorincz, P., Hegedus, K., Szeplaki, S., Low, P., & Juhasz, G. (2018). Molecular mechanisms of developmentally programmed crinophagy in Drosophila. J Cell Biol, 217(1), 361–374. doi:10.1083/jcb.201702145

Dittie, A. S., Hajibagheri, N., & Tooze, S. A. (1996). The AP-1 adaptor complex binds to immature secretory granules from PC12 cells, and is regulated by ADP-ribosylation factor. J Cell Biol, 132(4), 523–536. doi:10.1083/jcb.132.4.523

Gilleron, J., Gerdes, J. M., & Zeigerer, A. (2019). Metabolic regulation through the endosomal system. Traffic. doi:10.1111/tra.12670

Halban, P. A. (1991). Structural domains and molecular lifestyles of insulin and its precursors in the pancreatic beta cell. Diabetologia, 34(11), 767–778. doi:10.1007/bf00408349

Holst, B., Madsen, K. L., Jansen, A. M., Jin, C., Rickhag, M., Lund, V. K., … Gether, U. (2013). PICK1 deficiency impairs secretory vesicle biogenesis and leads to growth retardation and decreased glucose tolerance. PLoS Biol, 11(4) e1001542. doi:10.1371/journal.pbio.1001542 PBIOLOGY-D-12-04348 [pii]

Hou, J. C., Min, L., & Pessin, J. E. (2009). Insulin granule biogenesis, trafficking and exocytosis. Vitam Horm, 80, 473–506. doi:10.1016/S0083-6729(08)00616-X

Hummer, B. H., Maslar, D., Soltero-Gutierrez, M., de Leeuw, N. F., & Asensio, C. S. (2020). Differential sorting behavior for soluble and transmembrane cargoes at the trans-Golgi network in endocrine cells. Mol Biol Cell, 31(3), 157–166. doi:10.1091/mbc.E19-10-0561

Jiang, P., Nishimura, T., Sakamaki, Y., Itakura, E., Hatta, T., Natsume, T., & Mizushima, N. (2014). The HOPS complex mediates autophagosome-lysosome fusion through interaction with syntaxin 17. Mol Biol Cell, 25(8), 1327–1337. doi:10.1091/mbc.E13-08-0447

Keller, M. P., Choi, Y., Wang, P., Davis, D. B., Rabaglia, M. E., Oler, A. T., … Attie, A. D. (2008). A gene expression network model of type 2 diabetes links cell cycle regulation in islets with diabetes susceptibility. Genome Res, 18(5), 706–716. doi:10.1101/gr.074914.107

Klumperman, J., Kuliawat, R., Griffith, J. M., Geuze, H. J., & Arvan, P. (1998). Mannose 6-phosphate receptors are sorted from immature secretory granules via adaptor protein AP-I, clathrin, and syntaxin 6-positive vesicles. J Cell Biol, 141(2) 359–371. doi:10.1083/jcb.141.2.359

Kuliawat, R., Klumperman, J., Ludwig, T., & Arvan, P. (1997). Differential sorting of lysosomal enzymes out of the regulated secretory pathway in pancreatic beta-cells. J Cell Biol, 137(3), 595–608. doi:10.1083/jcb.137.3.595

Marsh, B. J., Soden, C., Alarcon, C., Wicksteed, B. L., Yaekura, K., Costin, A. J., … Rhodes, C. J. (2007). Regulated autophagy controls hormone content in secretory-deficient pancreatic endocrine beta-cells. Mol Endocrinol, 21(9), 2255–2269. doi:me.2007-0077 [pii] 10.1210/me.2007-0077

Ni, Q., Gu, Y., Xie, Y., Yin, Q., Zhang, H., Nie, A., … Wang, Q. (2017). Raptor regulates functional maturation of murine beta cells. Nat Commun, 8, 15755. doi:10.1038/ncomms15755

Orci, L., Ravazzola, M., Amherdt, M., Yanaihara, C., Yanaihara, N., Halban, P., … Perrelet, A. (1984). Insulin, not C-peptide (proinsulin), is present in crinophagic bodies of the pancreatic B-cell. J Cell Biol, 98(1), 222–228. Retrieved from https://www.ncbi.nlm.nih.gov/pubmed/6368567

Paquin, N., Murata, Y., Froehlich, A., Omura, D. T., Ailion, M., Pender, C. L., … Horvitz, H. R. (2016). The Conserved VPS-50 Protein Functions in Dense-Core Vesicle Maturation and Acidification and Controls Animal Behavior. Curr Biol, 26(7), 862–871. doi:10.1016/j.cub.2016.01.049

Park, J. J., Koshimizu, H., & Loh, Y. P. (2009). Biogenesis and transport of secretory granules to release site in neuroendocrine cells. J Mol Neurosci, 37(2), 151–159. doi:10.1007/s12031-008-9098-y

Pasquier, A., Vivot, K., Erbs, E., Spiegelhalter, C., Zhang, Z., Aubert, V., … Ricci, R. (2019). Lysosomal degradation of newly formed insulin granules contributes to beta cell failure in diabetes. Nat Commun, 10(1), 3312. doi:10.1038/s41467-019-11170-4

Pinheiro, P. S., Jansen, A. M., de Wit, H., Tawfik, B., Madsen, K. L., Verhage, M., … Sorensen, J. B. (2014). The BAR domain protein PICK1 controls vesicle number and size in adrenal chromaffin cells. J Neurosci, 34(32) 10688–10700. doi:10.1523/JNEUROSCI.5132-13.2014 34/32/10688 [pii]

Pols, M. S., van Meel, E., Oorschot, V., ten Brink, C., Fukuda, M., Swetha, M. G., … Klumperman, J. (2013). hVps41 and VAMP7 function in direct TGN to late endosome transport of lysosomal membrane proteins. Nat Commun, 4, 1361. doi:10.1038/ncomms2360

Price, A., Seals, D., Wickner, W., & Ungermann, C. (2000). The docking stage of yeast vacuole fusion requires the transfer of proteins from a cis-SNARE complex to a Rab/Ypt protein. J Cell Biol, 148(6), 1231–1238. doi:10.1083/jcb.148.6.1231

Riahi, Y., Wikstrom, J. D., Bachar-Wikstrom, E., Polin, N., Zucker, H., Lee, M. S., … Leibowitz, G. (2016). Autophagy is a major regulator of beta cell insulin homeostasis. Diabetologia, 59(7), 1480–1491. doi:10.1007/s00125-016-3868-9

Rieder, S. E., & Emr, S. D. (1997). A novel RING finger protein complex essential for a late step in protein transport to the yeast vacuole. Mol Biol Cell, 8(11), 2307–2327. doi:10.1091/mbc.8.11.2307

Sasidharan, N., Sumakovic, M., Hannemann, M., Hegermann, J., Liewald, J. F., Olendrowitz, C., … Eimer, S. (2012). RAB-5 and RAB-10 cooperate to regulate neuropeptide release in Caenorhabditis elegans. Proc Natl Acad Sci USA, 109(46), 18944–18949. doi:10.1073/pnas.1203306109 1203306109 [pii]

Seals, D. F., Eitzen, G., Margolis, N., Wickner, W. T., & Price, A. (2000). A Ypt/Rab effector complex containing the Sec1 homolog Vps33p is required for homotypic vacuole fusion. Proc Natl Acad Sci US A, 97(17), 9402–9407. doi:10.1073/pnas.97.17.9402

Sirkis, D. W., Edwards, R. H., & Asensio, C. S. (2013). Widespread Dysregulation of Peptide Hormone Release in Mice Lacking Adaptor Protein AP-3. PLoS Genet, 9(9), e1003812. doi:10.1371/journal.pgen.1003812 PGENETICS-D-13-00025 [pii]

Slot, J. W., & Geuze, H. J. (2007). Cryosectioning and immunolabeling. Nat Protoc, 2(10), 2480-2491. doi:10.1038/nprot.2007.365

Takats, S., Pircs, K., Nagy, P., Varga, A., Karpati, M., Hegedus, K., … Juhasz, G. (2014). Interaction of the HOPS complex with Syntaxin 17 mediates autophagosome clearance in Drosophila. Mol Biol Cell, 25(8), 1338–1354. doi:10.1091/mbc.E13-08-0449

Takeuchi, T., & Hosaka, M. (2008). Sorting mechanism of peptide hormones and biogenesis mechanism of secretory granules by secretogranin III, a cholesterol-binding protein, in endocrine cells. Curr Diabetes Rev, 4(1), 31–38. Retrieved from http://www.ncbi.nlm.nih.gov/pubmed/18220693

Thorens, B., Tarussio, D., Maestro, M. A., Rovira, M., Heikkila, E., & Ferrer, J. (2015). Ins1(Cre) knock-in mice for beta cell-specific gene recombination. Diabetologia, 58(3), 558–565. doi:10.1007/s00125-014-3468-5

Tooze, S. A. (1998). Biogenesis of secretory granules in the trans-Golgi network of neuroendocrine and endocrine cells. Biochim Biophys Acta, 1404(1-2), 231–244. doi:10.1016/s0167-4889(98)00059-7

Topalidou, I., Cattin-Ortola, J., Hummer, B., Asensio, C. S., & Ailion, M. (2020). EIPR1 controls dense-core vesicle cargo retention and EARP complex localization in insulin-secreting cells. Mol Biol Cell, 31(1), 59–79. doi:10.1091/mbc.E18-07-0469

Topalidou, I., Cattin-Ortola, J., Pappas, A. L., Cooper, K., Merrihew, G. E., MacCoss, M. J., & Ailion, M. (2016). The EARP Complex and Its Interactor EIPR-1 Are Required for Cargo Sorting to Dense-Core Vesicles. PLoS Genet, 12(5), e1006074. doi:10.1371/journal.pgen.1006074

Type_2_Diabetes_Knowledge_Portal. VPS41. type2diabetesgenetics.org. 2019 July 17; http://www.type2diabetesgenetics.org/gene/geneInfo/VPS41

van der Beek, J., Jonker, C., van der Welle, R., Liv, N., & Klumperman, J. (2019). CORVET, CHEVI and HOPS - multisubunit tethers of the endo-lysosomal system in health and disease. J Cell Sci, 132(10). doi:10.1242/jcs.189134

van der Welle, R. E. N., Jobling, R., Burns, C., Sanza, P., ten Brink, C., Fasano, A., … Klumperman, J. (2019). VPS41 recessive mutation causes ataxia and dystonia with retinal dystrophy and mental retardation by inhibiting HOPS function and mTORC1 signaling. bioRxiv, 2019.2012.2018.867333. doi:10.1101/2019.12.18.867333

Witwicka, H., Hwang, S. Y., Reyes-Gutierrez, P., Jia, H., Odgren, P. E., Donahue, L. R., … Odgren, P. R. (2015). Studies of OC-STAMP in Osteoclast Fusion: A New Knockout Mouse Model, Rescue of Cell Fusion, and Transmembrane Topology. PLoS One, 10(6), e0128275. doi:10.1371/journal.pone.0128275

Yamaguchi, H., Arakawa, S., Kanaseki, T., Miyatsuka, T., Fujitani, Y., Watada, H., … Shimizu, S. (2016). Golgi membrane-associated degradation pathway in yeast and mammals. EMBO J, 35(18), 1991–2007. doi:10.15252/embj.201593191

Zhang, X., Jiang, S., Mitok, K. A., Li, L., Attie, A. D., & Martin, T. F. J. (2017). BAIAP3, a C2 domain-containing Munc13 protein, controls the fate of dense-core vesicles in neuroendocrine cells. J Cell Biol, 216(7) 2151–2166. doi:10.1083/jcb.201702099

