## Supplemental Figures and Legends for "Pancreatic beta-cell specific deletion of VPS41 causes diabetes due to defects in insulin secretion"

**Supplemental Figures and Supplemental Figure Legends, Burns et al.**

**A**

Predicted Cleavage Site

WT INS-1 actctctcagactgagctATGG

VPS41 KO actctctcagactgagctATTGG
actctctcagactgagctAGTGG

**B**

VPS41 KO HA-VPS41

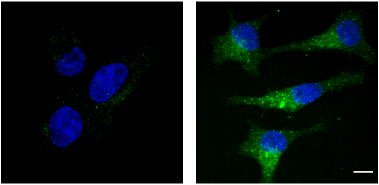

IF: HA

**Supplemental Figure 1 Burns et al.**

**Supplemental Figure 1**

(**A**) Predicted cleavage site for Cas9. Indels, both single base pair insertions, are shown in red in sequence disrupting the initiator ATG in Exon 1. (**B**) Representative images of VPS41 KO and HA-VPS41 rescue INS-1 cells immunostained for HA. Scale bar indicates 10µm.

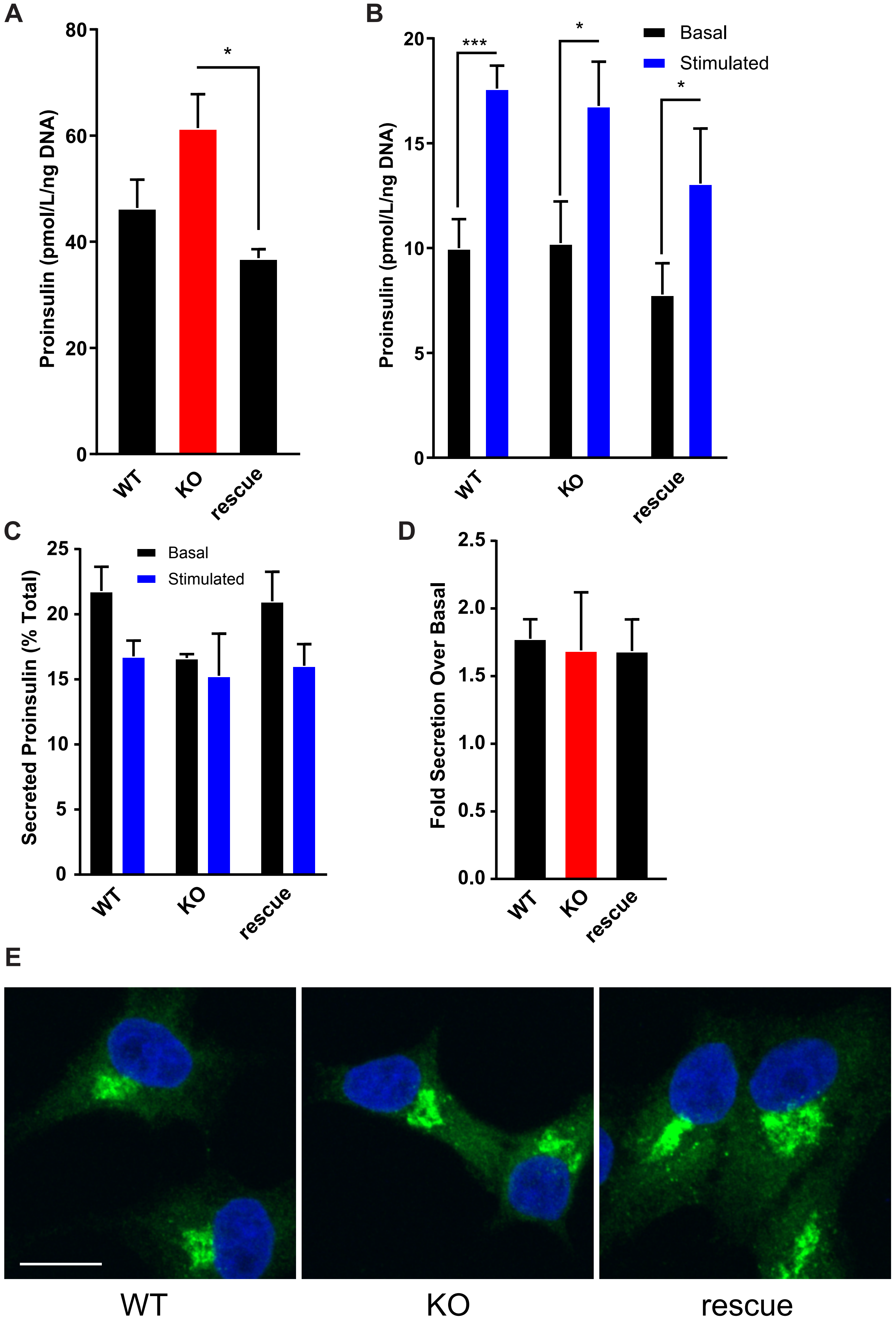


Supplemental Figure 2 Burns et al.

**Supplemental Figure 2**

(**A**) Total cellular proinsulin pool under basal conditions from VPS41 KO and rescue INS-1 was determined by ELISA. (**B**) Basal and stimulated proinsulin secreted from VPS41 KO and rescue INS-1 was determined by ELISA and analyzed by ANOVA. (**C**) Basal and stimulated proinsulin release from VPS41 KO and rescue INS-1 normalized to total cellular stores, analyzed by ANOVA. (**D**) Proinsulin secretion, fold stimulated release over basal release. Proinsulin cellular and secreted by ELISA. Data indicate mean +/- s.e.m.; n=3 independent experiments *: $p < 0.05$. ***: $p < 0.001$ analyzed by one way ANOVA.

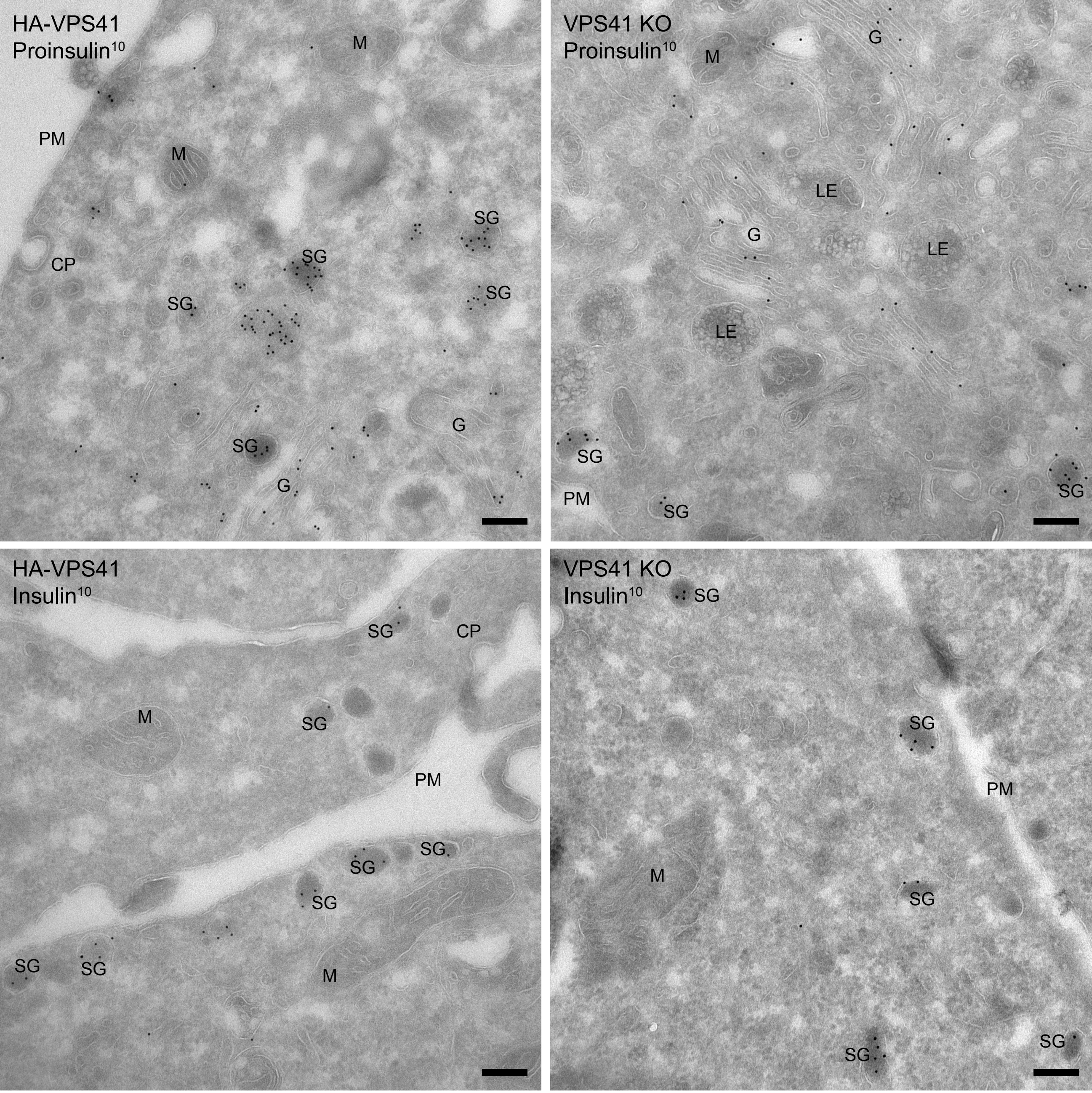


**Supplemental Figure 3 Burns et al.**

**Supplemental Figure 3**

Immuno-electron microscopy of VPS41KO or HA-VPS41 expressing INS-1 cells. Proinsulin (top) and insulin (bottom) localize to the lumen of secretory granules in both VPS41 KO and rescue cells expressing HA-VPS41 indicating that depletion of VPS41 does not result in missorting of (pro)insulin. CP= Coated pit, G= Golgi, LE= Late endosome, M= Mitochondria, PM= Plasma membrane, SG= Secretory granule. Scale bars: 200nm.

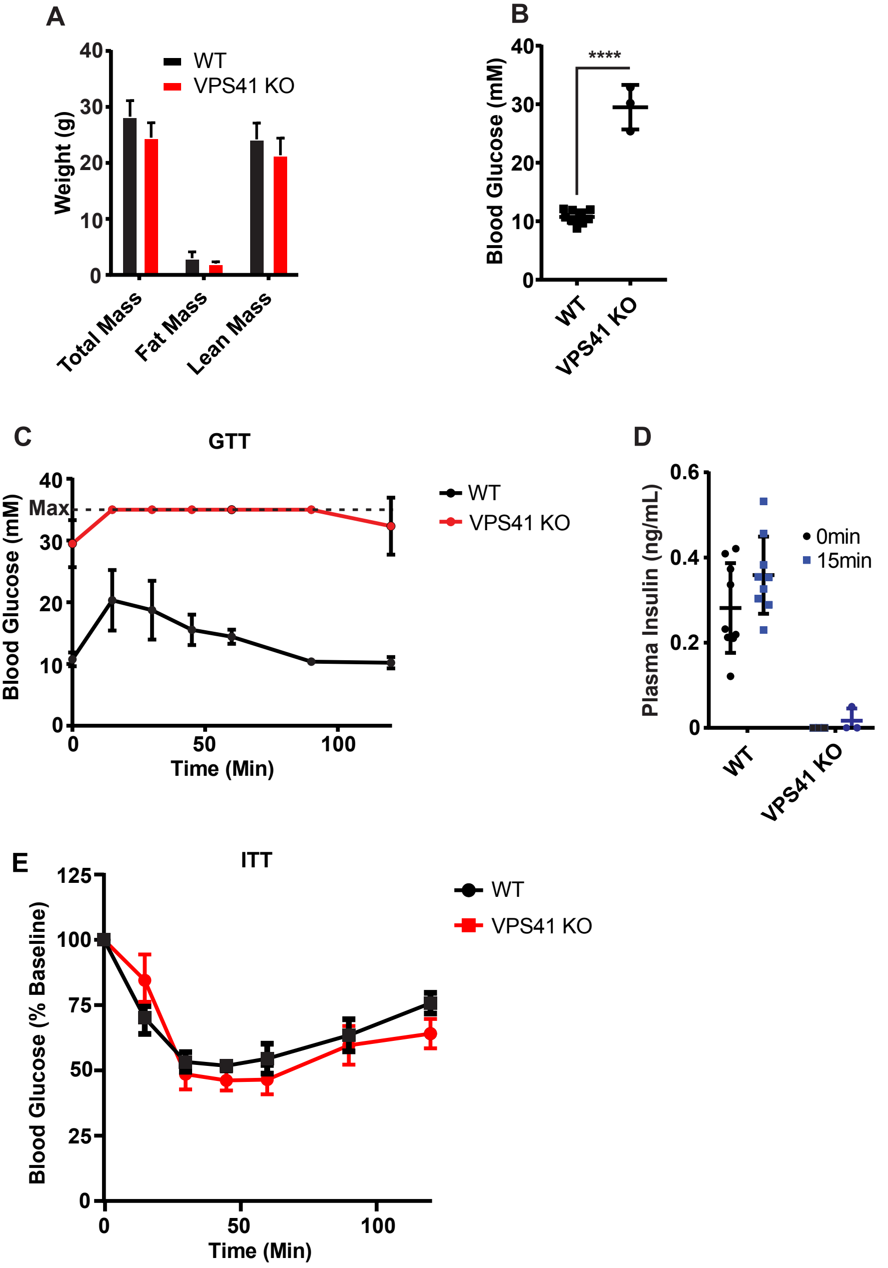


Supplemental Figure 4, Burns et al.

**Supplemental Figure 4**

(**A**) Fat and lean mass measurements of age matched 15-week-old WT and VPS41 KO mice. (**B**) Blood glucose measurement of mice fasted for 8hrs. (**C**) Blood glucose measurements during glucose tolerance test (GTT). The dotted lines indicate the maximum value of the glucometer. (**D**) Circulating blood insulin levels before and 15 min after glucose injection. Data indicate mean +/- s.e.m.; KO n=3, WT n=9 ****: $p < 0.0001$ secretion data analyzed by one-way ANOVA. (**E**) Blood glucose measurements during glucose tolerance test (ITT), values are normalized to the starting glycemia.

**Supplemental Movie 1**

3D array tomography of VPS41 KO INS-1 cells.

**Supplemental Movie 2**

3D array tomography of VPS41 KO INS-1 rescue cells.

**Supplemental Movie 3**

Representative movie of NPY-GFP RUSH in VPS41 KO cells. Scale bar indicated 10 µm.

**Supplemental Movie 4**

Representative movie of NPY-GFP RUSH in HA-VPS41 rescue cells. Scale bar indicated 10 µm.

**Supplemental Movie 5**

Representative movie showing basal and stimulated exocytosis of NPY-pHluorin in VPS41 KO INS-1 cells. Stimulation indicated by “High $K^+$”. Scale bar indicates 10 µm.

**Supplemental Movie 6**

Representative movie showing basal and stimulated exocytosis of NPY-pHluorin in HA-VPS41 rescue INS-1 cells. Stimulation indicated by “High $K^+$”. Scale bar indicates 10 µm.
